# Evolution rapidly optimizes stability and aggregation in lattice proteins despite pervasive landscape valleys and mazes

**DOI:** 10.1101/776450

**Authors:** Jason Bertram, Joanna Masel

## Abstract

Fitness landscapes are widely used to visualize the dynamics and long-term outcomes of evolution. The fitness landscapes of genetic sequences are characterized by high dimensionality and “ruggedness” due to sign epistasis. Ascending from low to high fitness on such landscapes can be difficult because adaptive trajectories get stuck at low-fitness local peaks. Compounding matters, recent computational complexity arguments have proposed that extremely long, winding adaptive paths may be required to even reach local peaks, a “maze-like” landscape topography. The extent to which peaks and mazes shape the mode and tempo of evolution is poorly understood due to empirical limitations and the abstractness of many landscape models. We develop a biophysically-grounded computational model of protein evolution based on two novel extensions of the classic hydrophobic-polar lattice model of protein folding. First, rather than just considering fold stability we account for the tradeoff between stability and aggregation propensity. Second, we use a “hydrophobic zipping” algorithm to kinetically generate ensembles of post-translationally folded structures. Our stability-aggregation fitness landscape exhibits extensive sign epistasis and local peaks galore. We confirm the postulated existence of maze-like topography in our biologically-grounded landscape. Although these landscape features frequently obstruct adaptive ascent to high fitness and virtually eliminate reproducibility of evolutionary outcomes, many adaptive paths do successfully complete the long ascent from low to high fitness. This delicate balance of “hard but possible” adaptation could occur more broadly provided that the optimal outcomes possible under a tradeoff are improved by rare constraint-breaking substitutions.

## 1 Introduction

The effect of an allele on organismal fitness can depend on its genetic context, a phenomenon known as epistasis. Epistasis is often visualized in terms of a “landscape” representing the mapping from high-dimensional genotype space to fitness (Wright, 1932; Kondrashov and Kondrashov, 2015). In the absence of epistasis, new mutations have the same effect regardless of the order in which they occur, and adaptive evolution on any genetic background leads to the same peak genotype(s). The latter still applies if epistasis only affects the magnitude — not the sign — of mutational fitness effects. However, if there is sign epistasis (mutations that are beneficial on some genetic backgrounds are deleterious on others), fitness landscapes become “rugged”; many or all of the mutational paths between two genotypes pass through valleys of lower fitness even if those genotypes only differ by a few mutations (Weinreich et al., 2005).

Since sign epistasis is pervasive, particularly at the level of individual proteins (Starr and Thornton, 2016), it is important to understand the consequences of landscape “ruggedness” (Kondrashov and Kondrashov, 2015). These consequences can be broadly be divided into two categories. First, sign epistatic landscapes can be permeated with local fitness peaks where no beneficial mutations are available, halting adaptive evolution and requiring valley crossing to proceed. Second, when sign epistasis occurs along realized adaptive paths, individual alleles alternate between being beneficial or deleterious as their genetic context evolves (Pollock et al., 2012; Shah et al., 2015; Starr et al., 2018), which can create a “maze-like” landscape on which adaptive evolution must backtrack by reverting alleles at some loci. In the extreme case that such reversions occur repeatedly across many loci, astronomically long adaptive paths may be required to reach even local fitness peaks (Kaznatcheev, 2019).

Although a large body of empirical work has explored the local properties of fitness landscapes (e.g. in the neighbourhood of some reference genotype) (De Visser and Krug, 2014; Starr and Thornton, 2016), the global properties of sign-epistatic fitness landscapes are still poorly understood (Kondrashov and Kondrashov, 2015). While globally mapping empirical landscapes is likely impossible, theoretical models should in principle allow global exploration (Fragata et al., 2018). One influential global approach is to analyze randomly constructed epistatic landscapes, but the applicability of these “random field models” (Stadler and Happel, 1999) to biological systems is unclear.

Theoretical toy models grounded in the biophysics of macromolecules hold more promise (Serohijos and Shakhnovich, 2014; Bastolla et al., 2017), but to date have been only applied in limited ways to study global landscape properties. The simplest approach for studying large macromolecular landscapes is to reduce fitness to a binary of “functional” and “non-functional” genotypes, creating a neutral network of functional genotypes connected by mutation (Maynard Smith, 1970). Global landscape structure then becomes a matter of evolutionary accessibility via neutral mutations among equivalently-high-fitness genotypes (Lipman and Wilbur, 1991; Wilke, 2001) (see Gavrilets (1999) for a similar approach applied at the organismal level). Even when this reduction to binary fitness is not made, many theoretical models focus on neutral landscape features such as mutational robustness (Van Nimwegen et al., 1999) and extended periods of neutral meandering (Fontana and Schuster, 1998; Ancel and Fontana, 2000). Consequently, far less is known about the global topography of fitness differences, even within the specific setting of macromolecules (Kondrashov and Kondrashov, 2015).

In particular, our understanding of how local peaks and mazes may impact evolution over long evolutionary trajectories — as might occur over the long-timescale global ascent from low to high fitness — is still rudimentary. Recent empirical studies suggest the existence of long-term trends in protein evolution, specifically an increase in hydrophobicity and a decrease in the clustering of hydrophobic amino acids along the primary sequence (Foy et al., 2019). The long timescale of these observed trends could conceivably be explained by slow valley-crossing processes (pervasive peaks) (Guo et al., 2019), or long adaptive paths (mazes), or a combination of these. Studies of macromolecular epistatic fitness landscapes with a global emphasis on long evolutionary paths should help to resolve such issues.

Here we investigate the global fitness landscape of an evolving protein using a computational lattice model to evaluate the mapping from amino acid sequence to molecular fitness. We evaluate thousands of adaptive trajectories to explore how difficult it is to reach high fitness, the diversity of high-fitness local peaks reached, the extent to which local peaks obstruct adaptation, and the prevalence of maze-like behavior. In response to the empirical results of Foy et al. (2019), we also explore whether our model is capable of generating long-term directional changes in sequence properties like hydrophobic clustering.

Following the classic approach of (Lau and Dill, 1989), our model groups amino acids as either hydrophobic (H) or polar (P) and maps protein structure onto a square lattice. Previous applications of HP lattice models to protein evolution have adopted a neutral network approach of focusing on “functional” amino acid sequences that have unique lowest-free-energy “native” conformations and evaluating whether such sequences are mutationally connected (Lipman and Wilbur, 1991; Chan and Bornberg-Bauer, 2002). Our approach differs in two respects.

First, we do not define fitness in terms of the existence of native conformations, which precludes all but the highest fitness HP sequences, and which requires extensive computational sampling of each sequence’s structural possibilities, necessitating the use of short sequences (of order ~ 20 residues). Instead, we implement a hydrophobic zipping algorithm that efficiently samples the molecular products of any sequence by simulating the kinetics of protein folding.

Second, along with fold stability, our protein fitness metric also accounts for the risk of inter-molecular aggregation. Aggregation risk is a crucial aspect of how proteins affect organismal fitness (DePristo et al., 2005; Levy et al., 2012), with important consequences for the fitness landscape of HP sequences. Since contacts between H residues are responsible for bonding both within (stable folding) and between (aggregation) protein molecules, there is a fundamental tradeoff between fold stability and aggregation potential. Innovation is made more difficult because P*→*H mutations will often cause both stability and aggregation potential to increase, and vice versa for H*→*P mutations. Importantly, the stability-aggregation tradeoff can induce sign epistasis as structural evolution changes each residue’s role. For instance, previously shielded H residues can become exposed and present an aggregation risk, while previously peripheral P residues selected for aggregation avoidance might become situated in more structurally important positions where an H residue would be better for stability. We find that evolution on our stability-aggregation fitness landscape is characterized by an abundance of local peaks and widespread maze-like behavior, causing strong path-dependence, largely eliminating reproducibility of outcomes at the sequence level, and frequently obstructing adaptive ascent to high fitness. Despite these obstacles, evolution does often manage to reach high fitness. Maze-like adaptive paths are created by pervasive sign epistasis, but are only of order *L* steps long, much shorter than the extremely long *O*(2^*L*^) paths that are hypothetically possible on sign-epistatic landscapes (Kaznatcheev, 2019). Adaptive paths that reach high fitness in our model can be seen as a series of innovations that cumulatively do better than naively expected from the intrinsic stability-aggregation tradeoff, a phenomenon that may apply more broadly wherever evolutionary tradeoffs are present.

## 2 Methods

### 2.1 Hydrophobic zipper model of protein stability and aggregation

We implement a two-dimensional HP lattice protein folding model. The amino acid heteropolymer is modeled as “beads on a string”, where each amino acid “bead” can be either hydrophobic (H) or polar (P). The allow-able conformations of the heteropolymer are self-avoiding walks on a two-dimensional square lattice (beads that are sequential neighbours must be one lattice step apart, and each lattice position holds at most one bead).

To be biologically useful, folded heteropolymers must be both thermo-dynamically stable and unlikely to aggregate with the other heteropolymers in the cellular environment. Following the classic HP model (Lau and Dill, 1989), the thermodynamic stability of a conformation (free energy of folding) is assumed to be proportional to the number of contacts between H monomers that are not sequential neighbours; this latter number we will refer to as “stability” *S*. Similarly, we define the “aggregation potential” *A* of a conformation as the number of potential H contacts that the conformation leaves exposed to bind to other molecules (Fig. 1). Combining these together, we assign an overall stability/aggregation score *F* = *S − A* to each conformation. The tradeoff between stability and aggregation (see Introduction) implies that *S* and *A* are strongly pleiotropic.

**Figure 1:**
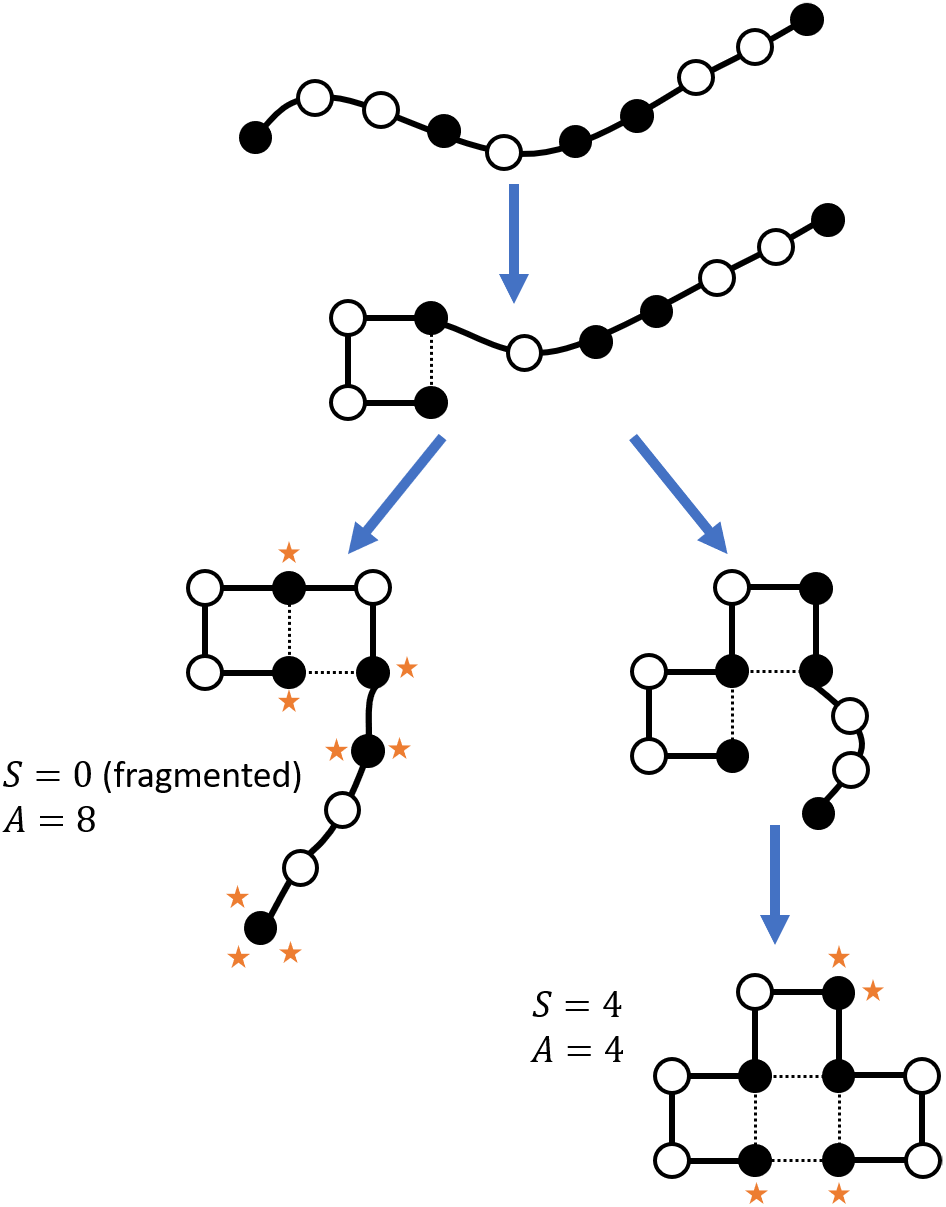
Hydrophobic zipping of the 10-monomer sequence HPPHPHHPPH (H: solid circles, P: hollow circles). Folding starts at the leftmost possible H-H contact (nucleation). Depending on the order in which H-H contacts form (a stochastic process), it is possible that the zipper gets “stuck” before much of the sequence is folded (left branch). Such sequences are regarded as fragmented and assigned *S* = 0. Orange stars denote potential H contacts contributing to the aggregation potential *A*. *S* is equal to the number of non-sequential H-H contacts (dotted lines).

We ensure that we are comparing the properties of sequences with the same length *L* as follows. It is not sufficient to simply fix the number of amino acids in the HP sequence, because sequences containing long runs of P residues are effectively collections of fragments shorter than *L* in the HP lattice zipper model. We therefore penalize fragmentation by setting *S* = 0 for zipped conformations that do not incorporate all *L* monomers in the sequence.

We model the kinetic process taking the amino acid heteropolymer from a disordered string to a folded globule as “hydrophobic zipping” (Dill et al., 1993). Our implementation follows the algorithm outlined in Dill and Fiebig (1994). The zipping process starts with an initial nucleation in which a pair of H monomers that are sequentially nearby — 3 amino acids apart in our square lattice model — form an H-H contact (Fig. 1). We assume that nucleation occurs between the leftmost such pair in the sequence (e.g. because folding begins before translation is complete; we return to this assumption in the Discussion). The formation of the nucleating contact brings other H monomers closer together, facilitating the formation of more H-H contacts. Each new H-H contact is chosen randomly from among the set of candidate H pairs that are possible given the current partially-zipped topology. To identify candidate H pairs, Dijkstra’s shortest path algorithm is applied to identify 3-step shortest paths connecting H residues on the graph of amino acid beads (nodes) connected by sequence-adjacency and existing H-H contacts (i.e. both solid and dotted lines in Fig. 1 are graph edges). A checking procedure excludes candidate pairs incompatible with the topology of the existing zipped structure. Zipping proceeds in this way until no further contacts are possible. Since fragmented sequences are assumed to have *S* = 0 in order to compare only sequences of comparable length, we do not allow further nucleation events after the initial one.

Hydrophobic zipping is a stochastic process because a given HP sequence *x* may zip to different conformations depending on the order in which H-H contacts form (Fig. 1). Consequently, *F*(*x*) is a random variable for each sequence *x*. We define the overall “fitness” 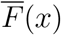 of *x* as the expectation of *F*(*x*). Although 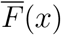 is not a direct measure of organismal or protein performance, and can even be negative, we use the term “fitness” for simplicity since 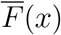 defines our fitness landscape. We estimate 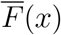 numerically by computing *F*(*x*) for a sample of 1000 zipped conformations and taking the sample mean.

Biophysically, hydrophobic zipping represents a rapid initial phase of conformational entropy loss (Dill et al., 1993). Zipping sometimes finds the lowest free energy conformations possible for a given sequence, but in general reaching the lowest free energy conformations requires breaking H-H contacts formed during zipping, a process that would presumably require longer timescales than the initial rapid collapse (Dill et al., 1993). Zipped conformations nevertheless approximate the lowest free energy conformations, and are produced with sample frequencies that reflect kinetic accessibility. With respect to aggregation, much of the aggregation risk associated with the expression of a sequence could be attributable to these rapidly-formed conformations.

### 2.2 Sequence evolution

We simulate sequence evolution as an origin-fixation process. Each substitutional step, we compute the fitnesses 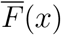 of the current sequence *x* as well as the fitnesses 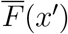 of all possible mutants *x*′ of *x*. We allow only single H *→* P or P *→* H mutations. There are *L* such mutants where *L* is sequence length. If fitter mutants are found to exist, one of them is chosen to replace *x*. We implement two alternative decision rules: choosing the fittest mutant and choosing a random fitter mutant. If all mutants are found to have lower fitness than *x*, we re-estimate 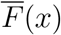 with a larger sample size and again check if the previously estimated mutant fitnesses exceed 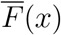. If no fitter mutants are found (a local peak), evolution is stopped; otherwise the above is repeated on the chosen fitter mutant. The re-estimation of 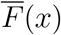 helps to ensure that we do not unintentionally miss any beneficial mutations due to sampling error in 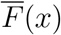.

We preclude mutations that create runs of three or more P monomers, since these would jam the zipper before all *L* amino acids could be incorporated in the fold. We also initialize evolution on sequences that start and end on H monomers and do not have runs of three or more P monomers, but are otherwise randomly generated. We do not specify initial hydrophobicity; instead, we vary the odds of H versus P monomers at each sequence position to generate a broad range of initial hydrophobicities.

### 2.3 Hydrophobic clustering

To connect our findings to the hydrophobic clustering results of Foy et al. (2019) we use the same hydrophobic clustering metric, defined as follows. Split an HP sequence of total length *L* into blocks of length *l*. In each block, subtract the number of P residues from the number of H residues to obtain a block score *n*. Hydrophobic clustering is then given by Ψ = *σ*^2^(*n*)/*K* where *σ*^2^(*n*) is the variance in *n* among blocks, and 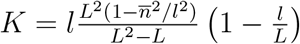 where 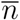 is the block mean *n* (Irbäck et al., 1996; Irbäck and Sandelin, 2000). Intuitively, Ψ is a modified dispersion index for the block scores *n* that accounts for finite sequence length *L* (similar to Bessel’s correction for estimating variance from a finite sample). The expectation of Ψ for completely random HP sequences with any *L* and any hydrophobocity is equal to 1 (Irbäck et al., 1996; Irbäck and Sandelin, 2000). We use a block length of *l* = 3 and report the average Ψ over the three possible block phases.

## 2.4 Data availability

Data and source code for simulations and figures can be accessed at https://github.com/jasonbertram/HPzipper_folding_vs_aggregation.

## 3 Results

## 3.1 Adaptive ascent from low-fitness random sequences to high fitness

In this section, we examine the global topography of the fitness landscape 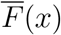 by simulating adaptive trajectories starting from different initial sequences generated randomly as described in Sec. 2.2. This gives a general sense of how likely it is to ascend to high fitness, and how adaptive evolution changes properties such as hydrophobicity and hydrophobic clustering. We assume a best-case scenario in which the most beneficial mutation is substituted at each step such that the fitness landscape is ascended at the greatest possible rate.

HP sequences can be divided into three qualitatively distinct groups related to their fitness 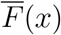. The highest fitness sequences always incorporate all *L* residues in their zipped conformations, and zip to only one or a few conformations (we call these sequences “complete”). The lowest fitness sequences never zip to a complete conformation (we call these sequences “incomplete”). “Intermediate” sequences only sometimes form a complete fold.

Our randomized initial sequences are almost always incomplete or intermediate, and have low fitness. The blue points in Fig. 2a-d show that adaptive evolution from these initial sequences frequently terminates at incomplete, low-fitness local peaks.

**Figure 2:**
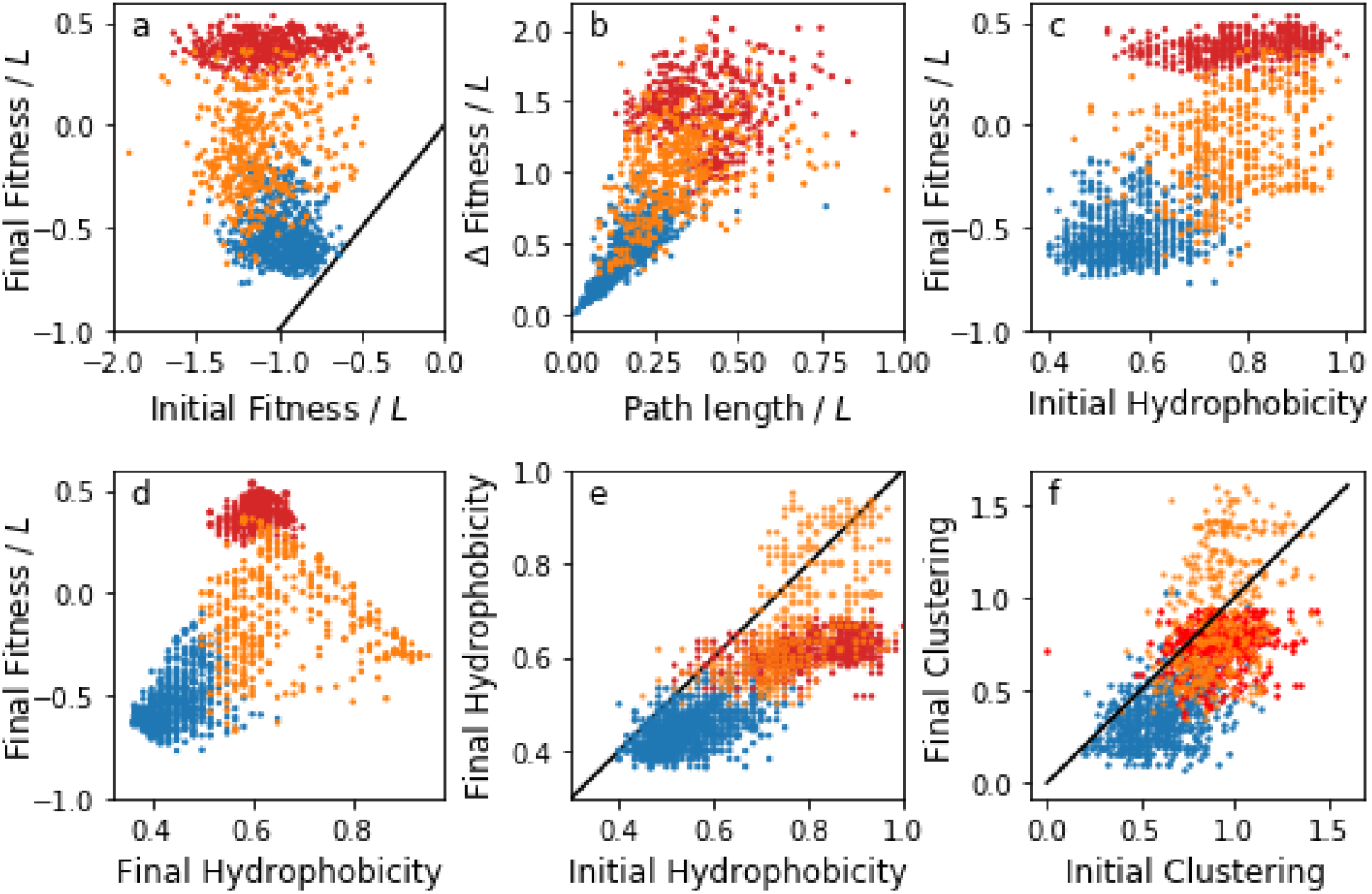
Adaptive origin-fixation evolution to a local peak on the fitness landscape 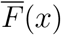 defined as stability minus aggregation following HP zipper folding. Complete sequences shown in red, incomplete sequences in blue, and intermediate sequences in orange. When sample points exactly overlap, the plotted circle is scaled proportional to the number of overlapping samples. Hydrophobic clustering uses a window size of 3 and averages over the three possible windowing phases. 1491 sequences of length *L* = 60 were evolved selecting the fittest beneficial mutation for each substitution.

Evolution does nevertheless manage to reach high-fitness complete sequences. Initial sequences which evolve completeness do not start with higher 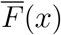 relative to other random initial sequences (Fig. 2a). Rather, it is the specific combination of H and P residues that determines whether high fitness is attainable. Local peaks that are complete sequences generally take more adaptive steps to reach than those with incomplete sequences, but there is considerable overlap in the adaptive path lengths (Fig. 2b). This means that some complete sequences experience long ascents in which beneficial mutations do not run out before reaching high fitness, whereas others attain high fitness via relatively quick shortcuts. Initial sequences with low hydrophobicity rarely reach a complete local peak, while initially high hydrophobicity improves the odds of reaching a complete sequence peak (Fig. 2c). However, many simulations with high initial hydrophobicity evolve to intermediate peaks short of completeness, indicating that reaching high fitness is also sensitive to the exact sequence of H and P residues.

Despite the sensitivity of fitness outcomes to genetic background, HP sequence evolution does exhibit consistent patterns. The highest fitness sequences all have hydrophobicity near *≈* 0.6 (Fig. 2d). Hydrophobicity tends to decline over an adaptive trajectory, although a significant minority of intermediate peak sequences gain H residues (Fig. 2e). Adaptive evolution also tends to reduce the within-sequence clustering of hydrophobic residues (Section 2.3).

### 3.2 Evolutionary mazes

In this section, we investigate the path-dependence of adaptive evolution on the stability-aggregation landscape 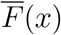 by repeatedly initializing evolution from the same sequence, but now picking a random beneficial mutation at each step rather than the highest fitness one as in the previous section. To ensure that it is possible to evolve completeness we choose a random initial sequence with a relatively high hydrophobicity of 0.78 (see Fig. 2c).

The landscape 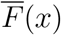 exhibits strong path-dependence. Despite starting from the same sequence, almost every adaptive path reached a different peak spanning incompleteness to completeness, depending on the order and identity of beneficial mutations (Fig. 3a). Adaptive paths also exhibit substantial “backtracking”, with total adaptive path length significantly exceeding the number of HP differences between the initial and final sequences (Fig. 3b). Taken together, these imply a maze-like landscape in which many poor path choices exist leading to low-fitness dead-ends, and meandering adaptive walks may be needed to reach high fitness due to alternation in the sign of mutational effects along adaptive trajectories (Fig. 4).

**Figure 3:**
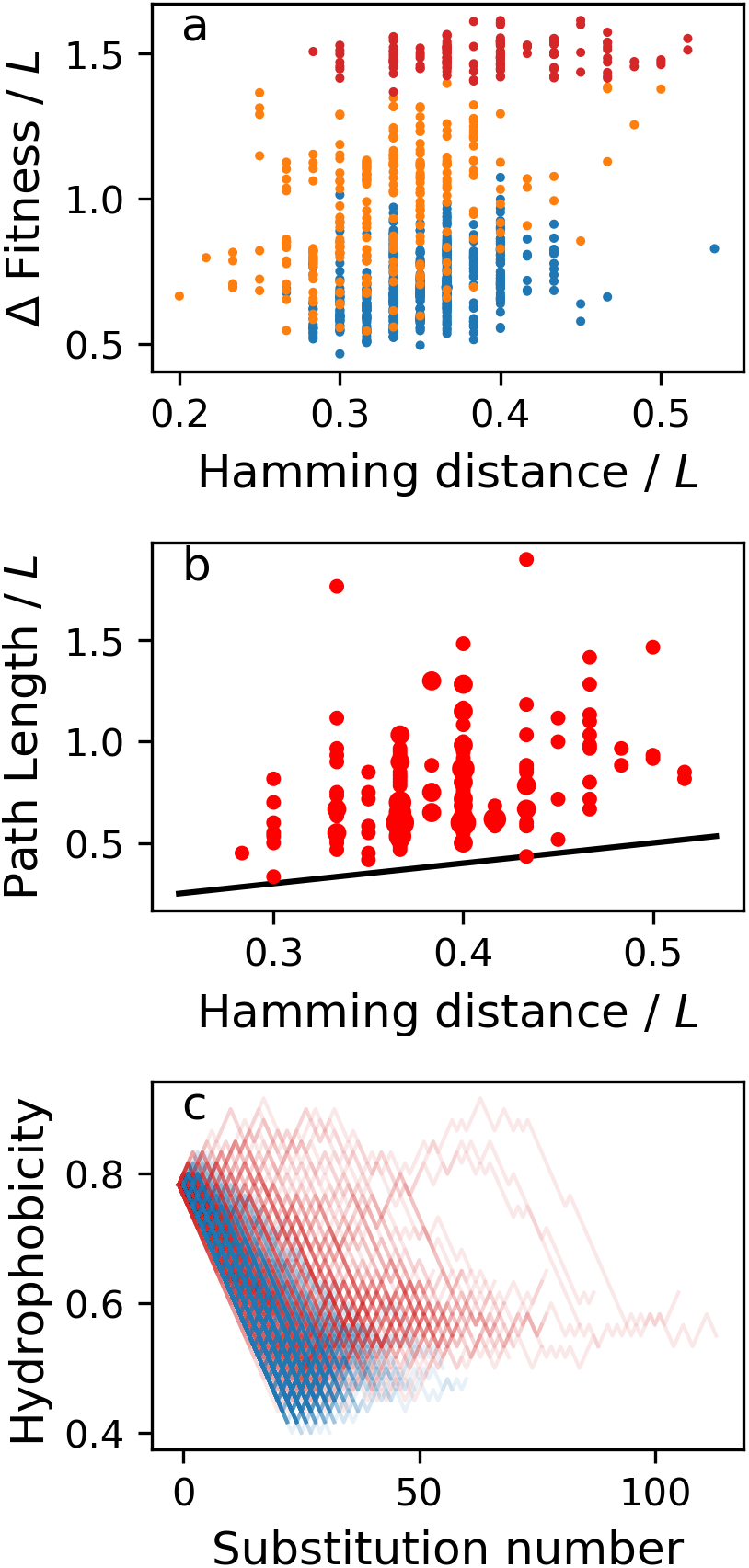
Maze-like behavior in HP lattice protein evolution. Starting from the same initial sequence, 987 sequences were evolved picking a random beneficial mutation at each step. (a) Many different evolutionary outcomes are possible. (b) Adaptive paths exhibit backtracking (only complete sequences shown for clarity; circle sizes proportional to number of sequences). (c) Evolution gets stuck at incomplete peaks (blue) when it focuses on removing hydrophobic residues to reduce aggregation propensity.

**Figure 4:**
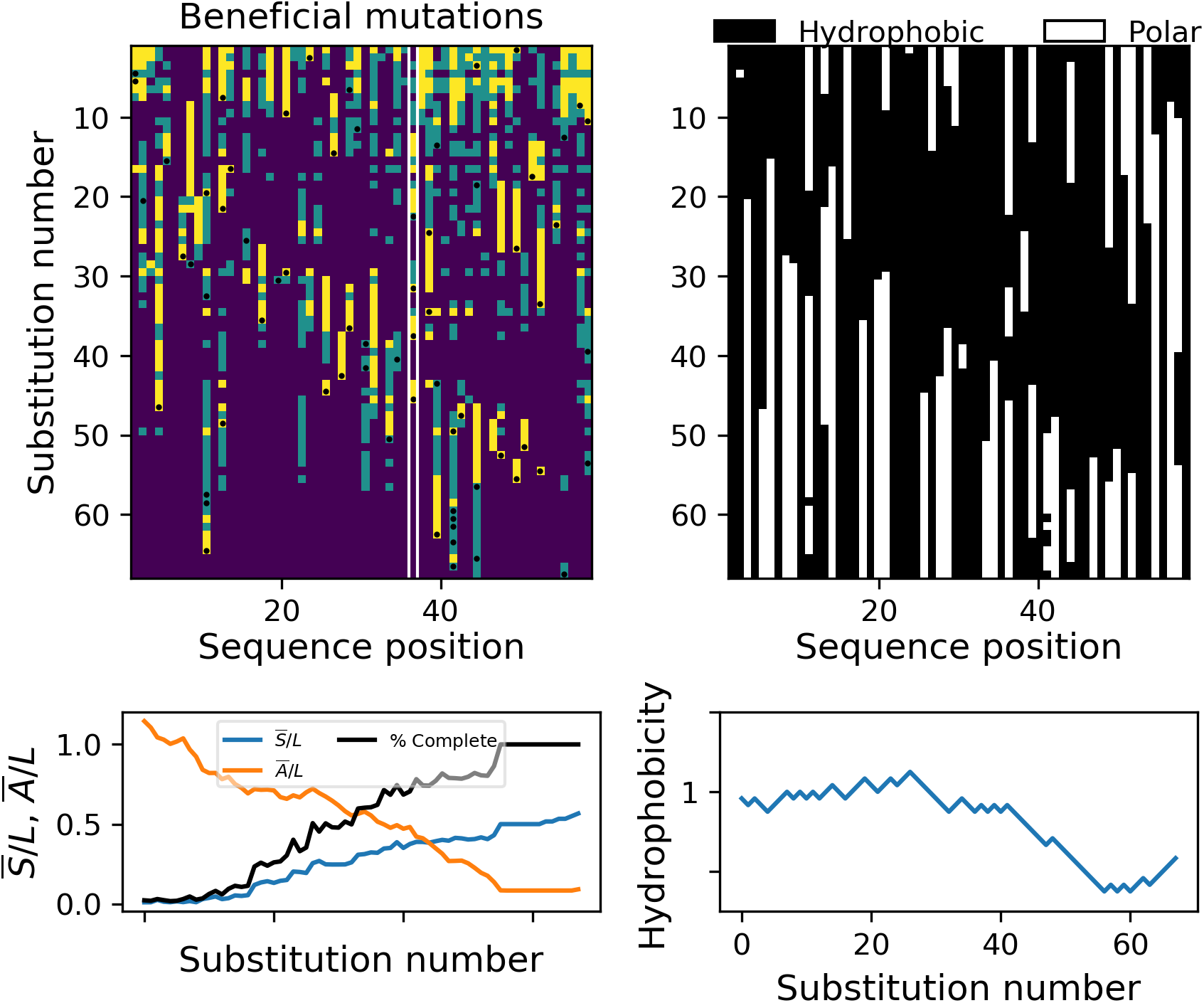
Example of a long (*> L*) adaptive path with backtracking. For the first *≈* 15 substitutions, evolution primarily reduces aggregation without reducing hydrophobicity. The chance of forming a complete fold then gradually increases, ultimately leading to completeness. Top left panel shows that the preferred state of most residues changes, in some cases repeatedly (highlighted with white lines). Beneficial mutations with 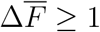 are yellow, deleterious mutations with 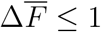 are dark and close-to-neutral mutations with 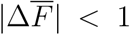 (whose scoring as beneficial/deleterious is more sensitive to sampling noise) are green. Black dots show mutations that occur (randomly chosen from among mutations with 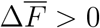). Residues with extended stretches of dark continuing to the peak have become “entrenched”. Evolving HP sequence on top right.

Adaptive paths that terminate at incomplete peaks are distinguished by the rapid removal of hydrophobic residues (Fig. 3c). This represents a lazy strategy of reducing aggregation risk by simply reducing overall hydrophobicity, dooming the prospects of finding complete folds. Adaptive paths that eventually attain completeness maintain H residues for longer, or even transiently accumulate them. In spite of this, in the early stages of adaptation, reductions in aggregation propensity *A* may still be the main source of gains in fitness *F* (Fig. 4). These paths therefore represent an ambitious strategy of tinkering with the fold to both reduce aggregation propensity and increase stability.

## 4 Discussion

We have shown that an HP lattice protein stability-aggregation fitness landscape exhibits pervasive sign epistasis, resulting in a high density of local peaks, evolutionary mazes, and strong path-dependence of evolutionary outcomes. One of the sources of sign epistasis in our model is that mutations can change the zipped conformations of a sequence in such a way that a residue that was buried in the protein core might now be exposed to the solvent. For an H residue, this will often mean switching from having a beneficial stabilizing role to posing a deleterious aggregation risk, and conversely for a P residue. Previous HP lattice models of protein evolution hinge on protein stability, not the intrinsic tradeoff between stability and aggregation and ways to optimize both despite the tradeoff, and as a result have not captured the full sign-switching repercussions of evolving different conformations (Lipman and Wilbur, 1991; Chan and Bornberg-Bauer, 2002; Bloom et al., 2006).

Despite the ruggedness of the stability-aggregation landscape, evolution from low-fitness random initial sequences frequently reaches high fitness without requiring valley crossing. These high-fitness adaptive paths may require back-and-forth H*↔*P mutations at some loci, and may consequently be longer than *L*. We therefore confirm, in a biologically-grounded model landscape, the importance of mazes posited by Kaznatcheev (2019) on the basis of more abstract landscape models. However, all of our adaptive paths are of order *L*, much shorter than the *O*(2^*L*^) paths that are hypothetically possible under sign epistasis. Intuitively, the latter exponential paths would require an amount of conformational rearrangement that is not consistent with gradually improving stability or aggregation potential at each adaptive step. Even along maze-like paths, some alleles become critical for maintaining accumulated fitness gains; further mutations at the corresponding loci are deleterious (right panel in Fig. 4). Such “entrenchment” (McCandlish et al., 2016; Shah et al., 2015; Ashenberg et al., 2013; Flynn et al., 2017; Starr et al., 2018) makes exponential-length adaptive paths unlikely, at least at the level of individual proteins.

One dimension in the fitness landscape — hydrophobicity — seems to strongly determine how far (Hamming distance), and how high (fitness), it is possible to move along adaptive paths. Complete evolved sequences tend to start at higher hydrophobicity (Fig. 2c) and maintain high hydrophobicity for longer (Fig. 3c) than incomplete sequences. Since the early stages in the evolution of a random sequence will generally entail high conformational diversity, many H residues will not have consistent structural roles and will therefore contribute little to expected stability 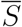. It is beneficial to replace these with P residues to reduce aggregation risk. However, since H residues are the raw material needed to fold, this “lazy” strategy closes off future avenues for innovation, potentially even committing evolution to further H residue removal. By contrast, transiently retaining or even adding H residues opens up conformational possibilities (a distinct mechanism from adding “surplus” stability to enable a stability-reducing innovation (Bloom et al., 2006)). This “industrious” strategy requires finding mutations that put the present H residues to better structural use.

Similar observations apply to hydrophobic clustering, which is strongly positively correlated with hydrophobicity both among initial sequences and following adaptive evolution. This correlation arises because, for most values of hydrophobicity, successful folding requires H residues to be more uniformly distributed than random along the sequence. For instance, among the randomly generated initial sequences, we require that no two H residues be more than three residues apart as this would surely cause zipping to fail (Section 2.2). Since lower hydrophobicity means fewer H residues to go around, lower hydrophobicity sequences will therefore tend to have lower clustering.

Remarkably, although evolutionary outcomes in our model are highly non-reproducible, even at the coarse level of fitness 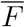 (Fig. 3), both hydrophobicity and hydrophobic clustering ultimately decline for almost all complete sequences (Fig. 2e,f). This appears to be a generic consequence of improving the consistency (chance to form a complete fold) of folding, since increases in hydrophobic clustering primarily occur in “intermediate” sequences which have high final hydrophobicity and, consequently, high conformational diversity. This may help to explain the long-term declines in hydrophobic clustering inferred empirically by Foy et al. (2019).

Our HP zipping model shares the usual disclaimer of all HP lattice models — though grossly simplifying reality, they nevertheless capture essential aspects of how protein folding is driven by contact between hydrophobic residues subject to conformational constraints (Dill and Fiebig, 1994). Our zipping model makes additional assumptions which add further caveats. It seems unlikely that our assumption of a single, fixed nucleation site applies to real proteins. In principle our findings could be sensitive to this assumption since it closes off adaptive paths whereby separately nucleated “subdomains” separated by disordered regions gradually combine into larger “complete” structures. However, we find that this scenario almost never occurs: in simulations where multiple nucleations are allowed, adaptive evolution almost always reaches a local peak consisting of a fragmented string of small (*~* 10 residues), highly-stable, independently nucleated structures. A related caveat is precluding H-H contacts from breaking once they have formed, which if allowed in a more sophisticated kinetic model might provide ways for separately nucleated substructures to merge. While we cannot rule out such possibilities, it seems plausible that the outcome of the rapid initial phase of folding representing by zipping is a good indicator of the stability and aggregation properties of a sequence.

The stability-aggregation tradeoff creates an evolutionary quandary in which mutations that increase (decrease) a positive attribute (stability) often also increase (decrease) a negative attribute (aggregation propensity). Beneficial mutations will consequently often only constitute a small proportion of all possible mutations (Fig. 4). Moreover, as with any evolutionary tradeoff, some of these beneficial mutations only improve the balance between the two sides of the tradeoff (e.g. by gaining more in stability than aggregation potential). Given these constraints on adaptation, how are long adaptive trajectories leading from globally-low to globally-high fitness possible? The answer is that the quandary between stability and aggregation is not a strict tradeoff: there exist (rare) mutations that are able to escape the status quo or even represent a “win-win” (both gaining stability and lowering aggregation potential).

It seems plausible that fitness landscapes with similar “improvable trade-off” properties could occur more widely than the protein folding landscape studied here. The stability-aggregation tradeoff in our model is an example of “frustration” caused by competing interactions (Wolf et al., 2018). Frustration seems to be a universal feature of biological macromolecules (Ferreiro et al., 2014, 2018), where competition occurs between the tendency to engage in beneficial interactions and the tendency to engage in deleterious interactions, i.e. between affinity and specificity. Competition between affinity and specificity also constrains evolution at the cellular level of protein-protein interaction networks; similar to our findings, evolutionary adaptation can still occur by adjusting the fitness consequences of this inescapable conflict Heo et al. (2011). More broadly, frustration effects occur across scales of biological organisation, as evidenced by the ubiquity of tradeoffs (Wolf et al., 2018). The prevalence of constraint-breaking mutations is less apparent, but environmental change, which is also ubiquitous, can act as a transient constraint-breaker (De Vos et al., 2015). Our protein fitness landscape model can thus be viewed as a case study of the evolutionary effects of frustration, with lessons that may extend well beyond protein evolution.

## Acknowledgements

This work was supported by the John Templeton Foundation (60814). We thank David McCandlish for comments on the manuscript, and Artem Kaznatcheev for discussions. The authors acknowledge the Indiana University Pervasive Technology Institute for providing HPC Karst resources that have contributed to the research results reported within this paper. This research was supported in part by Lilly Endowment, Inc., through its support for the Indiana University Pervasive Technology Institute, and in part by the Indiana METACyt Initiative. The Indiana METACyt Initiative at IU was also supported in part by Lilly Endowment, Inc.

